# The ERA-related GTPase AtERG2 associated with mitochondria 18S RNA is essential for early embryo development in *Arabidopsis*

**DOI:** 10.1101/190173

**Authors:** Pengyu Cheng, Hongjuan Li, Linlin Yuan, Huiyong Li, Lele Xi, Junjie Zhang, Jin Liu, Yingdian Wang, Heping Zhao, Huixin Zhao, Shengcheng Han

## Abstract

The ERA (*E. coli* RAS-like protein)-related GTPase (ERG) is a nuclear-encoded GTPase with two conserved domains: a GTPase domain and a K Homology domain. ERG plays a vital role in early seed development in *Antirrhinum majus*. However, the mechanism that regulates seed development remains unclear. Blasting the genome sequence revealed two homologies of ERG, AtERG1, and AtERG2 in *Arabidopsis*. In this study, we found that AtERG2 is localised in the mitochondria and binds mitochondrial 18S RNA. Promoter and transcript analyses indicated that *AtERG2* was mainly expressed in the leaf vein, trichome, mature pollen, and ovule. The mutants of *AtERG2* showed recessive lethal, gametophytic maternal effects, silique shortage, and early seed abortion, in which some seeds arrested in the zygotic stage at 1.5 days after pollination (DAP) and aborted at 2.0 DAP in *aterg2-1* +/-. Reactive oxygen species (ROS) accumulated at 1.5 DAP in the arrested seeds, and the transcription of several ROS-responsible genes, *WRKY40*, *ANAC017*, and *AOX1a*, was up-regulated in the *aterg2-1* +/- seeds which were arrested 1.5 and 2.0 DAP but not in wild-type (WT) and *aterg2-1* +/- seeds. The cell death-related gene BAG6 was also transcriptionally activated in *aterg2-1* +/- seeds arrested at 2.0 DAP. Chloramphenicol treatment during pollination induced a similar phenotype and gene expression pattern but showed no transcriptional changes of ANAC017 in WT. These results suggested that *AtERG2* promotes early seed development by affecting the maturation of the mitochondria ribosome small subunit and mitochondrial protein translation in *Arabidopsis*.

## Introduction

The small GTP-binding proteins (GTPases) are found in all domains of life and act as molecular switches that are “activated” by GTP and “inactivated” by the hydrolysis of GTP to GDP to regulate numerous cellular processes, such as signal transduction, reorganisation of the cellular cytoskeleton, regulation of transcription and translation, protein transport and vesicle trafficking (Takai et al., 2001; Vernoud et al., 2003; Shan, 2016). Based on the sequences and structural signatures, the small GTPase superclass can be divided into two large classes: the <u>tra</u>nslation <u>fac</u>tors (TRAFAC) class and the <u>si</u>gnal recognition particle (SRP), <u>Mi</u>nD-like ATPases, and <u>Bi</u>oD (SIMIBI) class. The TRAFAC class includes proteins involved in translation, signal transduction (in particular, the extended Ras-like family), and cell motility. The SIMIBI class is involved in protein localisation, chromosome partitioning, membrane transport, and a group of metabolic enzymes with kinase or related phosphate transferase activity (Leipe et al., 2002). In 1986, a yeast Ras-like protein that contains 316 amino acids and low GTP hydrolyse activity was found in *E.coli* and named *E.coli* Ras-like protein (ERA) (Ahnn et al., 1986). ERA is essential for cell growth and viability (Inada et al., 1989), cell division (Britton et al., 1997), and pleiotropic processes, including carbon metabolism, fatty acid metabolism, and adaptation to thermal stress (Lerner and Inouye, 1991; Voshol et al., 2015). ERA has two conserved domains: the GTPase domain, which is homologous to RAS and belongs to the TRAFAC class, and the K Homology (KH) domain, which binds to the *E.coli* 16S rRNA and the 30S ribosomal subunits (Meier et al., 1999). Tu *et al*. showed that the GTP-binding form of ERA is helpful for recognition and binding to the (1530)GAUCACCUCC(1539) sequence at the 3' end of 16S rRNA, and RNA recognition stimulates its GTP-hydrolysing activity and the switch to the GDP-binding form; the latter suggested that ERA acts as a chaperone for the processing and maturation of 16S rRNA to benefit the assembly of the 30S ribosomal subunit (Tu et al., 2009; Tu et al., 2011).

ERA is highly conserved among bacteria, archaea, and eukaryotes, and its homologue in humans is called Era-like 1 (ERAL1), which is nuclear-encoded and localised in the mitochondria matrix (Britton et al., 2000; Uchiumi et al., 2010). Previous studies showed that ERAL1 binds mitochondria 12S rRNA, but not cytoplasmic ribosome rRNA, to help the maturation of the mitochondria small ribosomal subunit, which is similar to Era (Dennerlein et al., 2010; Uchiumi et al., 2010; Uchiumi and Kang, 2012). Prior studies showed that knockdown of ERAL1 by siRNA inhibits mitochondrial protein translation and elevates mitochondrial reactive oxygen species (ROS) production, leading to cell death (Uchiumi et al., 2010; Xie et al., 2012). These results indicated that ERAL1 plays an important role in the small ribosomal constitution and is involved in cell viability.

In plants, *ERA-related GTPase* (ERG) was first found in *Antirrhinum majus* and is expressed in dividing or metabolically active cells. ERG was predicted to target to the mitochondria by amino-terminal sequences, and a deletion allele of *ERG* by site-selected transposon mutagenesis resulted in the arrested development of seeds containing embryos and endosperm after fertilisation (Ingram et al., 1998). Genomic data on the *ERG* genes in *Arabidopsis* showed that there are two homologues, AtERG1, localised in chloroplast, and AtERG2, localised in the mitochondria (Suwastika et al., 2014). Jeon *et al* (Jeon et al., 2014) showed that *Nicotiana benthamiana* double Era-like GTPase (NbDER) contains two tandemly repeated GTPase domains and a C-terminal KH-like domain involved in RNA binding. NbDER was localised primarily to chloroplast nucleoids, possessed GTPase activity, and bound to 23S and 16S ribosomal RNAs, which contributed to the maturation and assembly of the chloroplast 50S ribosomal subunit. Depletion of NbDER impaired processing of plastid-encoded ribosomal RNAs, resulting in accumulation of the precursor rRNAs in the chloroplasts and a leaf-yellowing phenotype caused by disrupted chloroplast biogenesis. However, the mechanism of AtERGs involved in the organelle rRNA processing and ribosome biogenesis remains unclear. Here, we showed that AtERG2 was localised in the mitochondria, dependent on its N-terminal sequence, and bound mitochondria 18S RNA *in vivo*. The T-DNA insertion mutant of *ATERG2* is recessive lethal, and some seeds arrest at the zygote stage at 1.5 DAP and abort at 2.0 DAP in *aterg2-1* +/-. We further found that ROS were accumulated, and the transcription of several ROS-responsible genes, WRKY40, ANAC017, and AOX1a, and the cell death-related gene, BAG6, were up-regulated in the *aterg2-1* +/- arrested seeds but not in WT and *aterg2-1* +/- developed seeds; this is similar to CAP treatment with WT. These results suggest that AtERG2 is involved in regulating seed early development through mitochondria ribosome maturation and protein translation.

## Materials and Methods

### Plant growth and chloramphenicol treatment

*Arabidopsis thaliana* (Columbia-0) plants were grown in soil with a 16/8-h light/dark cycle at 22°C in the plant growth room. Seeds were sterilised and plated onto 1/2 Murashige and Skoog (MS) basal salt medium with 0.8% agar and 1.5% (w/v) sucrose in the dark for 2 days at 4°C. Then, the plates were grown in the growth chamber (Panasonic, Japan) at a 16/8-h light/dark cycle at 22°C for 7 days, and the seedlings were transplanted in the soil for continual growth. One week after stemming, the pre-pollinated flowers were treated by dipping the whole flower into 500 μg /mL Chloramphenicol (CAP) in 0.02% Sliwet-77 solution for 30 s. The treatment lasted for 3 days, and the siliques were collected at the exact developmental stages for the next analysis. The control was the same buffer without CAP.

### DNA and RNA extraction

*Arabidopsis* genomic DNA was isolated from 4-week-old leaves using the extraction buffer (200 mM Tris-HCl pH 7.5, 250 mM NaCl, 25 mM EDTA, 0.5% SDS) as described (Edwards et al., 1991). Total RNA was extracted by TRIZOL Reagent (Thermo Fisher Scientific, USA) from *Arabidopsis* seeds according to the manufacturer’s instructions (Thermo Fisher Scientific) and quantified using a Nanodrop ND-1000 spectrophotometer (LabTech, USA). Total RNA (5 μg) was used for cDNA synthesis with Superscript II RNase H-Reverse Transcriptase (Life Technologies, USA) and oligo(dT)18 primers according to the manufacturer’s instructions.

### Characterisation of T-DNA Insertion Mutants

Two T-DNA insertion lines of AtERG2: *aterg2-1* (SALK_032115) and *aterg2-2* (SALK_032124) were purchased from *Arabidopsis* stock centres (http://www.arabidopsis.org/) (Figure 3A). The genotype of T-DNA lines was confirmed by PCR (Figure S3A) using the gene-specific primers and a T-DNA–specific primer (Table S1).

### Constructs and transgenic plants

The coding sequence of *AtERG2* was amplified by PCR and cloned into the Gateway Cloning vector pE6c or pE3c (Addgene, USA) with Bam HI and Not I sites. To generate the various truncated *AtERG2* constructs, such as AtERG2_N (containing only N-terminal), AtERG2_C (containing GTPase domain (GD) and KH domain (KH)), AtERG2_GD (containing only GD), and AtERG2_KH (containing only KH), the corresponding AtERG2 fragments were amplified and subcloned into pE6c. Construct maps containing different parts of *AtERG2* are shown in Figure S2. The different constructs were used in combination with the destination vector pMDC32 (http://www.arabidopsis.org/) with the Gateway LR II kit (Life Technologies, USA) to generate the plant expression vectors. A genomic fragment of 1.8 kb in the promoter region of *AtERG2* was amplified by PCR and cloned into a GUS binary vector pCAMBIA 1305.1 with EcoR I and Bgl II sites. The primers used in these constructs are listed in Table S1. All vectors were confirmed by sequencing.

The resulting 35S::AtERG2-6XMyc and P_*AtERG2*_::GUS constructs were transformed into *Agrobacterium tumefaciens* GV3101. Transgenic plants were generated by the floral dip method (Clough and Bent, 1998) and selected on MS medium containing hygromycin B (25 mg/L) or kanamycin (50 mg/L). More than 100 AtERG2-6XMyc or *P_AtERG2_*::GUS lines were recovered. Independent homozygous transformants carrying a single insertion in the T3 generation were further analysed.

### Transient transformation of *Arabidopsis* mesophyll protoplasts

The vectors containing AtERG2-eYFPs were co-transformed with a mCherry-tagged organelle marker gene and transiently expressed in *Arabidopsis* mesophyll protoplasts, as described previously (Liang et al., 2015). After incubation for 16 h at 22°C, fluorescence was visualised using an LSM 700 confocal microscope (Zeiss, Germany). Observations were made using a 63× objective under oil immersion. eYFP fluorescence was excited at 488 nm and collected at SP 550 IR. The mCherry fluorescence was stimulated at 555 nm and collected at SP 630 IR.

### Immunoprecipitation of mitochondria 18S RNA with AtERG2

The 2-week-old transgenic *Arabidopsis* seedlings overexpressing AtERG2-6Xmyc were ground in liquid nitrogen, and the protein/RNA complexes were extracted using equal volumes of lysis buffer (IP buffer (50 mM NaCl, 2 mM MgCl_2_, 10 mM HEPES pH 7.5, 2 mM DTT, 2 mM EDTA, 100 U/mL RNasin RNase inhibitor [Promega, USA], and 100 U/mL protease inhibitor cocktail [Sigma-Aldrich, USA] plus 1% TritonX-100 and 0.5% SDS) and incubated for 1 h on ice, then centrifuged twice at 13,000 g for 10 min at 4°C to remove the insoluble debris. The supernatant was diluted with nine volumes of IP Buffer and incubated with 10 μL anti-Myc antibody (Sigma-Aldrich) for 4 h on ice with occasional gentle mixing. Next, the anti-Myc-decorated extracts were incubated with 10 μl protein G agarose beads pre-washed with IP buffer for 1 h at 4°C with rotation. Then, the beads were spun down at 10,000 rpm in a microfuge for 1 min at 4°C, and washed eight times with 1 mL washing buffer (100 mM NaCl, 2 mM MgCl_2_, 10 mM HEPES pH 7.5, 2 mM DTT, 2 mM EDTA, 100 U/mL RNasin RNase inhibitor, and 100 U/mL protease inhibitor cocktail). After elution with TRIZOL reagent, co-immunoprecipitated RNA was isolated and analysed by quantitative real-time RT-PCR (qRT-PCR).

### qRT-PCR

qRT-PCR was performed on the 7500 Fast Real-Time PCR System (Applied Biosystems, USA) using Power SYBR^®^ Green PCR Master Mix (Applied Biosystems). The thermal program was 2 min at 50°C, 10 min at 95°C, followed by 40 cycles of 15 s at 95°C and 60 s at 60°C. The data were normalised to the expression of *Arabidopsis actin*. The dissociation curve program was used to confirm the specificity of the target amplification product. All primers used in this study are listed in Table S1. At least three independent biological replicates were performed for qRT-PCR analysis. The expression analysis was performed using the Data Processing System. A one-way analysis of variance (ANOVA) and Tukey’s multiple range test were conducted to determine significant differences. P < 0.05 was considered statistically significant.

### Microscopy analysis

Seeds were removed from siliques at different developmental stages, cleared in Hoyers solution (7.5 g gum arabic, 100 g chloral hydrate, 5 mL glycerol in 30 mL water) and examined using a Zeiss Axio Observer A1 microscope equipped with DIC optics as described by Liu and Meinke (Liu and Meinke, 1998). The ROS detection was performed according to a modified version of a previously described method (Daudi et al., 2012) by staining with 1 mg/mL 3,3'-diaminobenzidine (DAB, Sigma) containing Triton X-100 (0.1% v/v) and 10 mM sodium phosphate buffer (pH 7.0). As previously described (Yu et al., 2005), the GUS staining was performed for 4 hours with seedlings and leaves or overnight with flowers and seeds at 37°C in GUS staining solution (50 mM sodium phosphate buffer, pH 7.0, 10 mM EDTA, 0.1% Triton X-100, 2 mM potassium ferricyanide, 2 mM potassium ferrocyanide, 1 mg/mL X-Gluc) before the clearing procedure.

## Results

### Phylogenetic relationship of AtERGs and its homologues

A previous study showed that there are two homologues of ERG in the *Arabidopsis* genome: AtERG1 (At5g66470), localised in the chloroplast, and AtERG2 (At1g30960), localised in the mitochondria (Suwastika et al., 2014). We used the sequence of ERA, ERAL1, ERG, AtERG1, and AtERG2 to search NCBI public databases (http://blast.ncbi.nlm.nih.gov) at an E-value of 1e-10. The presence of conserved GTPase and KH domains is the exclusive criterion for confirmation of 16 ERA homologous proteins in 14 species, which include MmERA from *Mus musculus*, DrERA from *Danio rerio*, DmERA from *Drosophila melanogaster*; ScERA from *Saccharomyces cerevisiae*, OsERG1 and OsERG2 from *Oryza sativa*, CsERG from *Cucumis sativus*, ZmERG from *Zea mays*, NtERG from *Nicotiana tabacum*, StERG from *Solanum tuberosum*, and SlERG from *Solanum lycopersicum*. Additionally, Jeon *et al*. (Jeon et al., 2014) found a chloroplast-localised double Era-like GTPase in *Nicotiana benthamiana* (NbDER), which binds to chloroplast 23S and 16S ribosomal RNAs. We further identified the homologues of NbDER, such as AtDER from *Arabidopsis thaliana*, EcDER from *E. coli*, ZmDER from *Zea mays*, and OsDER from *Oryza sativa*. Then, a phylogenetic tree was generated using the CDS (sequence coding for amino acids in protein) of 21 ERA homologues, and we found that members of the ERA homologues were separated into four distinct clades, designated I, II, III, and IV (Figure S1). Clade I included 12 members: ERG, AtERG2, CsERG, OsERG2, ZmERG, NtERG, SIERG, StERG, ERAL1, MmERAL1, DrERAL1, and DmERAL1, which were predicted and/or identified as mitochondria-localised. Clade II contained three members: ERA, AtERG1, and OsERG1; this suggested that ERA is most closely related to chloroplast-localised AtERG1 and OsERG1. Clade III included one member: ScERA. Clade IV had five DERs. These eukaryotic ERALs and ERGs were nuclear-encoded and organelle-localised genes, indicating that they were transferred to the nucleus after organelle symbiogenesis.

### AtERG2 is dependent on its N-terminal sequence for localisation to the mitochondria

To further investigate the subcellular localisation of AtERG2, we developed various constructs containing different parts of AtERG2 fused with yellow fluorescent protein (eYFP) at their C-terminal end (Figure S2) and transiently co-transformed cells with a mCherry-labelled organelle marker in the *Arabidopsis* mesophyll protoplasts. The full-length AtERG2 was localised in the mitochondria as previously reported (Figure 1A) (Suwastika et al., 2014). We further found that the N-terminal end (1-150 aa) of AtERG2 was localised in the mitochondria. However, the C-terminal end, GTPase domain, and KH domain alone diffused into the cytoplasm (Figure 1A). These results indicated that the N-terminal end of AtERG2 is critical for its localisation to the mitochondria.

**Figure 1.**
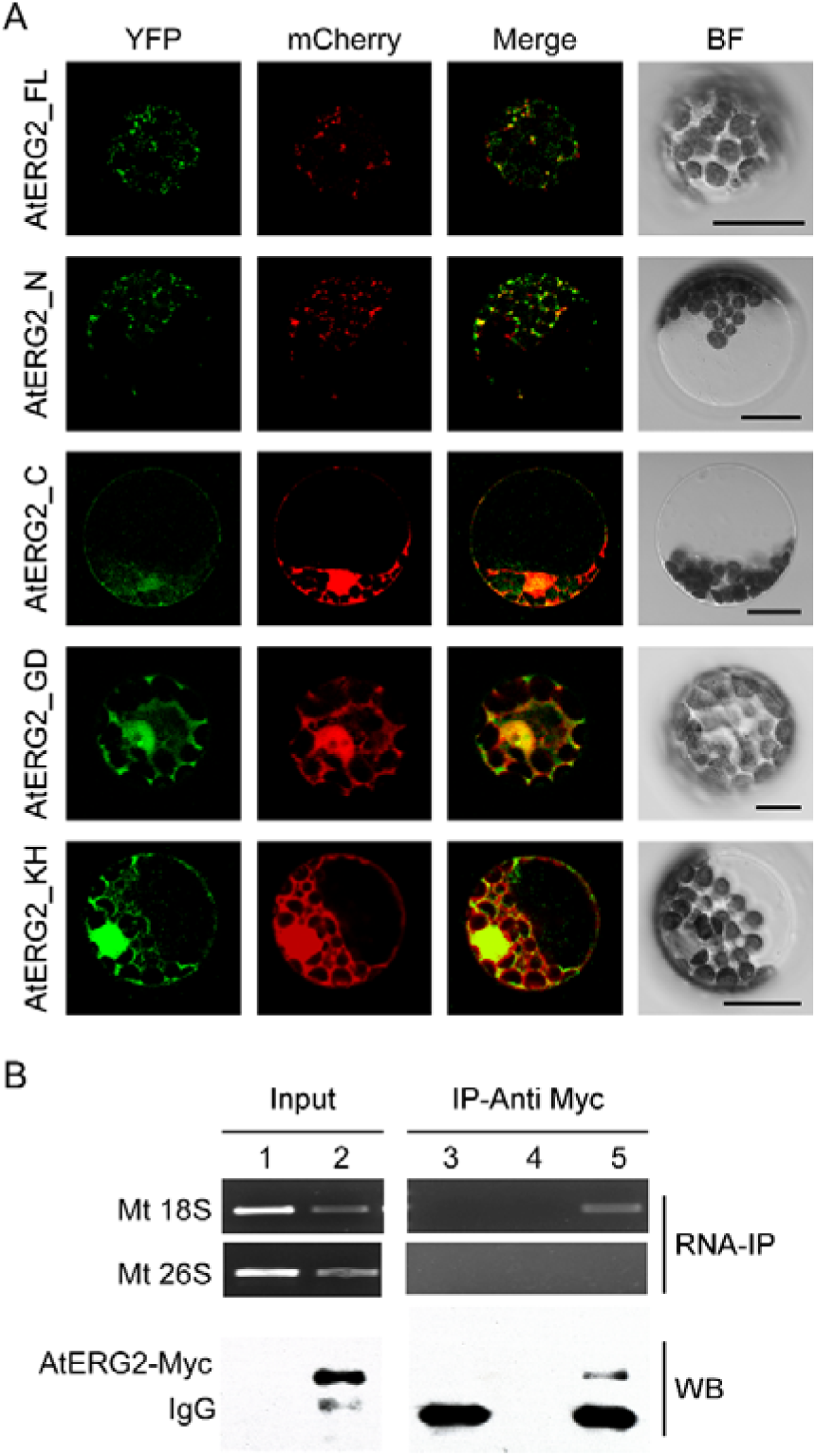
AtERG2 is localised in mitochondria and associated with mitochondria 18S RNA. (A) Localisation of various truncated AtERG2 constructs in *Arabidopsis* mesophyll cells. The full-length AtERG2 (AtERG2_FL-YFP) and AtERG2 N-terminal part (AtERG2_N-YFP) were co-transformed with mCherry-labelled mitochondria marker; AtERG2 C-part (AtERG2_C-YFP), AtERG2 GTPase domain (AtERG2_GD-YFP), and AtERG2 KH domain (AtERG2_KH-YFP) co-transformed with mCherry-labelled cytoplasm marker. All images in this figure were obtained from one optic section. Scale bars are equivalent to 20 m. The schematic of different AtERG2-eYFP constructs were shown in Figure S3. (B) RNA immunoprecipitation (RNA-IP) assays (top) showing the interaction between AtERG2-MYC and Mt 18S RNA in *Arabidopsis* were performed with (Lane 3 and Lane 5) or without (Lane 4) MYC antibody followed by RT-PCR detection. Mt 28S RNA served as a negative control (Lane 1 and Lane 2). The Western blot (WB, bottom) indicates the protein level in the transgenic plants, which was detected prior to the RNA-IP assay.

### AtERG2 associates with mitochondria 18S rRNA *in vivo*

We next used RNA immunoprecipitation (RNA-IP) assays to detect AtERG2 binding ribosome rRNA. The AtERG2-Myc:rRNA complex was extracted from the transgenic *Arabidopsis* stably expressed AtERG2-Myc fusion protein, preincubated with Myc antibody, and purified by protein G agarose beads. Then, the rRNA was isolated with Trizol solution and detected by RT-PCR (Figure 1B). The results showed that AtERG2 binds mitochondria ribosome small subunit 18S rRNA but not large subunit component 26S rRNA, suggesting that AtERG2 plays the same role as its homologues ERA and ERAL1 for the maturation of 16S rRNA and assembly of ribosomes.

### *AtERG2* is mainly expressed in the leaf vein, trichome, mature pollen, and ovule

To detect the transcription pattern of AtERG2 in *Arabidopsis*, we first generated the independent transgenic *Arabidopsis* lines, which stably expressed *P*_*AtERG2*_::GUS (Figure 2). In the F2 generation, GUS activity was observed in the leaf vein of 1-week-old seedling cytoledons (Figure 2A), trichomes of 3-week-old rosette leaves (Figure 2B and 2C), pollen grains inside the anthers (Figure 2D and 2E), and ovules during the period of pollination and early-stage seed development (Figure 2F-I). In anthers, GUS activity was relatively weak the day before pollination (Figure 2D), but it increased in mature pollen at 0.5 DAP (Figure 2E). We also found that GUS staining was detected at a low level in the pre-matured embryo sac before pollination (Figure 2F); it then reached the highest level in the ovule of 0.5- and 1.0-DAP seeds during the zygotic stage (Figure 2G and 2H) and quickly dropped in the 1.5-DAP seeds as soon as it reached the early-globe stage (Figure 2I). These results were further confirmed by qRT-PCR assays to evaluate the expression of AtERG2 in different tissues and in the development of *Arabidopsis*. Figure 2J showed that AtERG2 transcripts were highly expressed in the 0.5-, 1.0-, and 1.5-DAP seeds.

**Figure 2.**
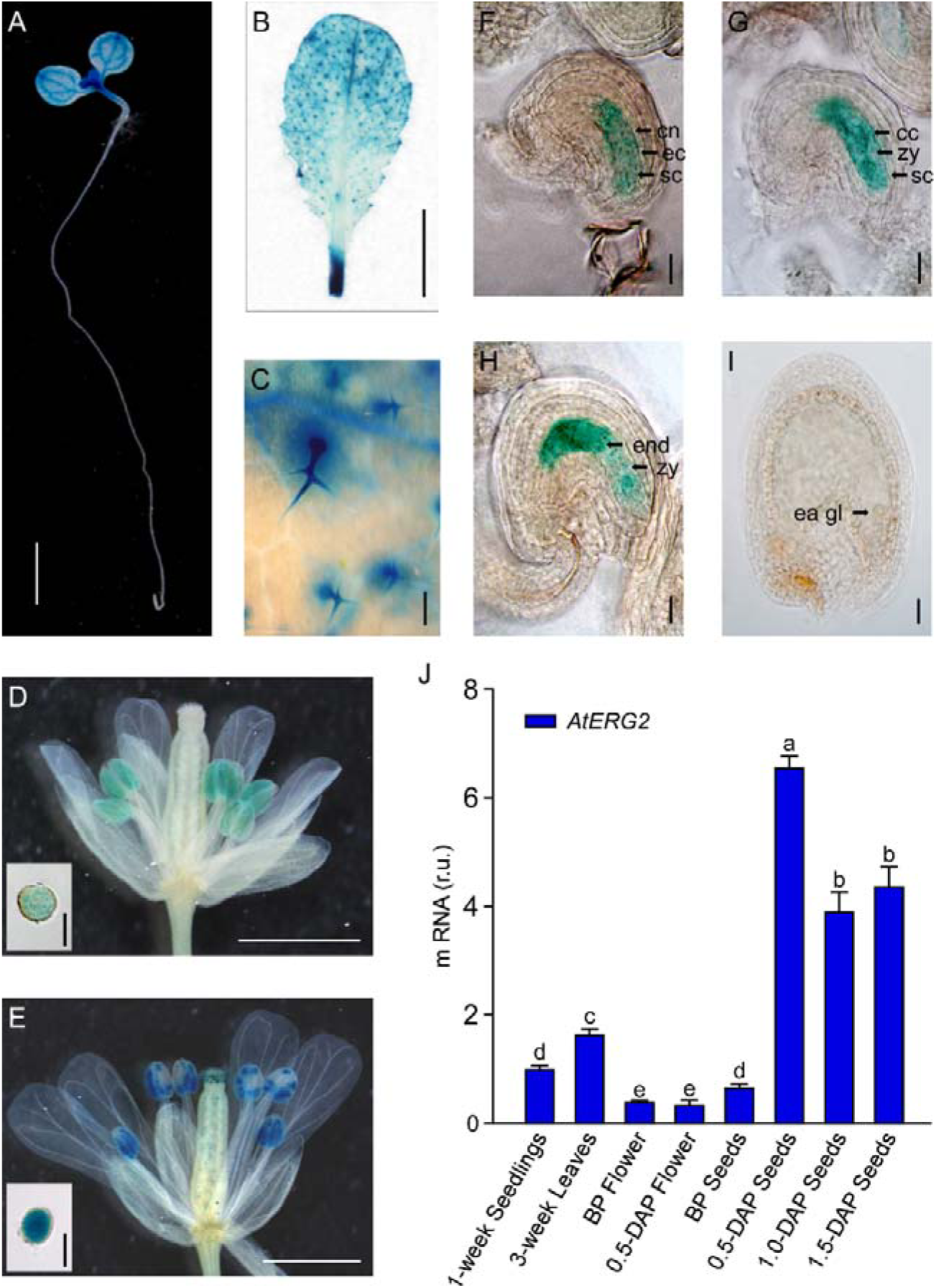
Differential Expression of *AtERG2* in pollen and early-stage ovule. GUS staining of P_*AtERG2*_·:GUS transgenic *Arabidopsis* plants for 4 hours with seedlings and leaves or overnight with flowers and seeds. Bar = 2 mm in (A); 5 mm in (B); 0.2 mm in (C); 0.5 mm in (D) and (E); 0.05 mm in (F), (G), (H), and (I); and 0.02 mm in pollen of (D) and (E). (A) *P* _*AÆRG2*_::GUS fusion staining in 1-week-old *Arabidopsis* seedling showing GUS activity in cotyledon and shoot apical meristem. (B) and (C) *P*_*AtERG2*:_:GUS fusion staining in 4-week-old *Arabidopsis* leaves and leaf trichome. (D) and (E) *PAtERG2*:: GUS fusion staining in flowers and pollen at BP (before pollination) and 0.5 DAP (day after pollination), respectively. (F) to (I) *PAtERG2*::GUS fusion staining in developed seeds at BP, 0.5, 1.0, and 1.5 DAP, respectively. The central polar nuclei are indicated by cn; the egg cells are indicated by ec; the synergid cells are indicated by sc; the central triploid nucleus is indicated by cc; the endosperm are indicated by end; the embryos at zygote are indicated by zy; and the early globe-stage embryos (with fewer than four cells in a single embryo) are indicated by ea gl. (J) The transcription level of *AtERG2* is detected by qRT-PCR in different tissues of *Arabidopsis*. UBQ10 was used as internal control. Error bars indicate the SD of three biological replicates. Different letters (a–e) indicate statistically significant differences (p < 0.05) according to Tukey’s multiple range test.

### The T-DNA insertion mutant of *AtERG2* shows a recessive lethal and gametophytic maternal effect (GME) phenotype

To characterise the function of AtERG2, the two T-DNA insertion lines (SALK_032115, named as *aterg2-1*, and SALK_032124, named as *aterg2-2*, respectively, Figure 3A) were obtained from The *Arabidopsis* Information Resource (TAIR, https://www.arabidopsis.org/), and the genotype of the progeny from self-crossed plants was examined by PCR. There were no homozygous mutant seeds from the progeny of self-crossed *aterg2-1* or *aterg2-2* plants (Figure S3A), and the segregation ratio of WT to *aterg2-1* +/- was close to 1:1, which indicated that the mutant line was recessive lethal (Table 1). Then, we backcrossed *aterg2-1* +/- with WT plants and found that the segregation ratio of WT to *aterg2-1* +/- was 2:1 when using WT as a paternal provider, and the ratio of WT to *aterg2-1* +/- was almost 1:1 when using WT as the maternal provider (Table 1). These results showed that *aterg2-1* has a GME phenotype, indicating that AtERG2 plays a vital role in regulating *Arabidopsis* seed development.

**Figure 3.**
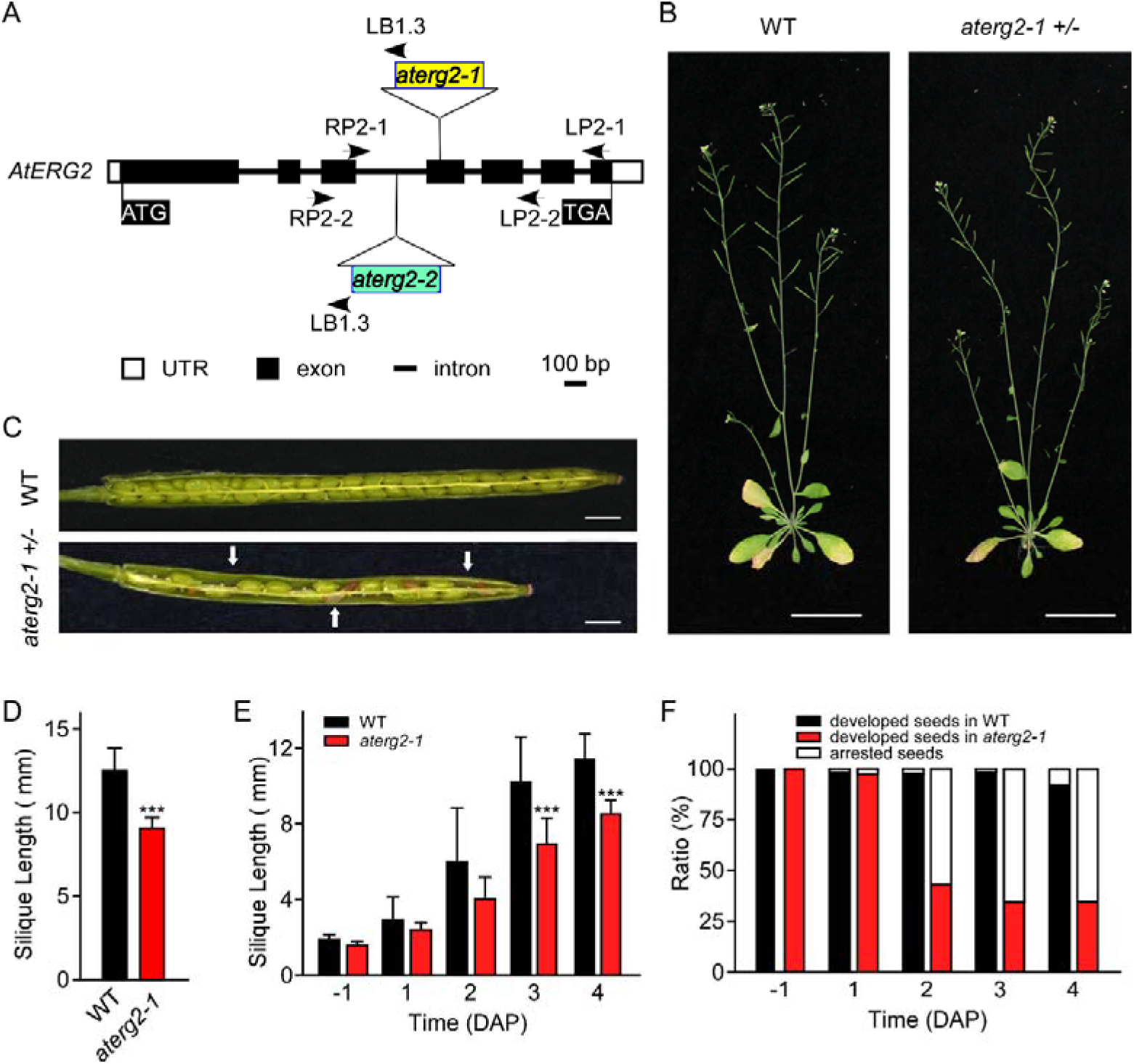
*AtERG2* is essential for early seedling development. (A) The schematic diagram for two T-DNA insertions in the *Arabidopsis* genome of *aterg2*. Two lines, SALK_032115 (*aterg2-1*) and SALK_032124 (*aterg2-2*), were purchased from The *Arabidopsis* Information Resource (TAIR, http://www.arabidopsis.org/). Open box, UTR; black box, exon; line, intron. Bar = 100 bp. (B) The 6-week-old whole aerial parts of the wild-type (WT) and heterozygous mutant *aterg2-1* +/- after planting in soil. Bar = 5 cm. (C) The opened 6-DAP (day after pollination) siliques from WT and *aterg2-1* +/-. The arrows point to the aborted seeds. Bar = 1 mm. (D) The length of mature siliques from WT and *aterg2-1* +/- (n = 60). The values are means ± SD. *** indicates extremely statistically significant values (P < 0.01). (E) The silique length of WT and *aterg2-1* +/- measured from the day before pollination to four DAP (n = 16). The values are means ± SD. *** indicates extremely statistically significant values (P < 0.01). (F) The seed abortion ratio of WT and *aterg2-1* +/- calculated from the day before pollination to 4 DAP (silique number = 8).

**Table 1.** Genotype of the *aterg2* +/- mutant

| Parental Genotype (Female x Male)<sup>a</sup> | *AtERG2* | *aterg2* +/- | *aterg2* | ratio<sup>b</sup> |
| --- | --- | --- | --- | --- |
| *aterg2* +/- × *aterg2* +/- | 153 | 166 | 0 | 0.921 : 1 |
| *aterg2* +/- × *AtERG2* | 21 | 10 |  | 2.1 : 1 |
| *AtERG2* × *aterg2* +/- | 47 | 36 |  | 1.306 : 1 |

<sup>a</sup> Plants were crossed manually, and seeds of the resulting cross were collected and grown on MS plates. The germinated seedlings were transferred to soil, and their 4-week-old leaves were then collected for genomic DNA extraction. Their genotypes were determined by PCR.
<sup>b</sup> The ratio is the number of plants exhibiting the WT genome: the number of plants with the *aterg2* +/- genome without the T-DNA insertion.

### *aterg2-1* +/- shows defects in early embryo development

During the whole growth period from germination to flowering, both *aterg2-1* and *aterg2-2* showed similar phenotypes to WT, with the exception of silique length and seed number; also, the ratios of the developed seeds of mutant lines were lower than those of the WT (Figure 3B to 3D and Figure S3B and S3C). We found that the developed seeds per silique in *aterg2-1* and *aterg2-2* were about 30% of WT at 6 DAP until seed maturation (Figure S3B and S3C), which suggested that seed development was arrested and aborted at the early development of seeds in both *aterg2-1* and *aterg2-2*. Therefore, we used *aterg2-1* +/- to perform the following experiments.

Next, we investigated the silique development of WT and *aterg2-1* +/- from the pre-pollination stage (BP) to 4 days after pollination (DAPs) and found that the silique length was significantly shorter at 3 DAP in *aterg2-1* +/- than in WT (Figure 3E). Some seeds were small and arrested at 1 DAP, and the arrested ratio of seed development reached 65% at 3 DAP in *aterg2-1* +/- (Figure 3F), which is almost the same as the ratio of seed development at 6 DAP. We further dissected the siliques and hyalinised seeds to observe the embryo development at BP, 0.5, 1.5 and 2.0 DAP to explore the impairment of early seed development in *aterg2-1* +/-. Firstly, we found that there is no developmental difference of embryo sac between WT and *aterg2-1* +/- plants at BP and 0.5 DAP (Figure S4). In addition, we showed that there are 50.7% and 64.2% small-sized seeds at 1.5 DAP and 2.0 DAP in *aterg2-1* +/- silique, respectively, but only 2.4% were found in WT silique, although the silique length was the same between WT and *aterg2-1* +/- plants at 2.0 DAP (Figure 4A and 4B). At 1.5 DAP, there were two types of embryos in WT and *aterg2-1* +/- siliques: 31.1% in the zygote stage and 68.9% in the early-globe stage for WT and 56.9% in the zygote stage and 43.1% in the early globe-stage embryos for *aterg2-1* +/- siliques, respectivley (Figure 4C). Furthermore, we found 9.8% heart-stage, 57.9% globe-stage (eight or more cells of embryos), and 32.2% early globe-stage embryos in 2.0-DAP WT seeds, and 1.3% heat-stage, 21.1% globe-stage, and 23.2% early globe-stage embryos were found in 2.0-DAP *aterg2-1* +/- seeds. More interestingly, as much as 38% of seeds stayed in the zygote stage, and 16.5% of these seeds were collapsed seeds without embryos in 2.0-DAP *aterg2-1* +/- siliques (Figure 4C). These results suggested that insertional inactivation of AtERG2 causes delay in the development of embryos after pollination and arrested seeds from 1.5 DAP in *aterg2-1* +/- plants.

**Figure 4.**
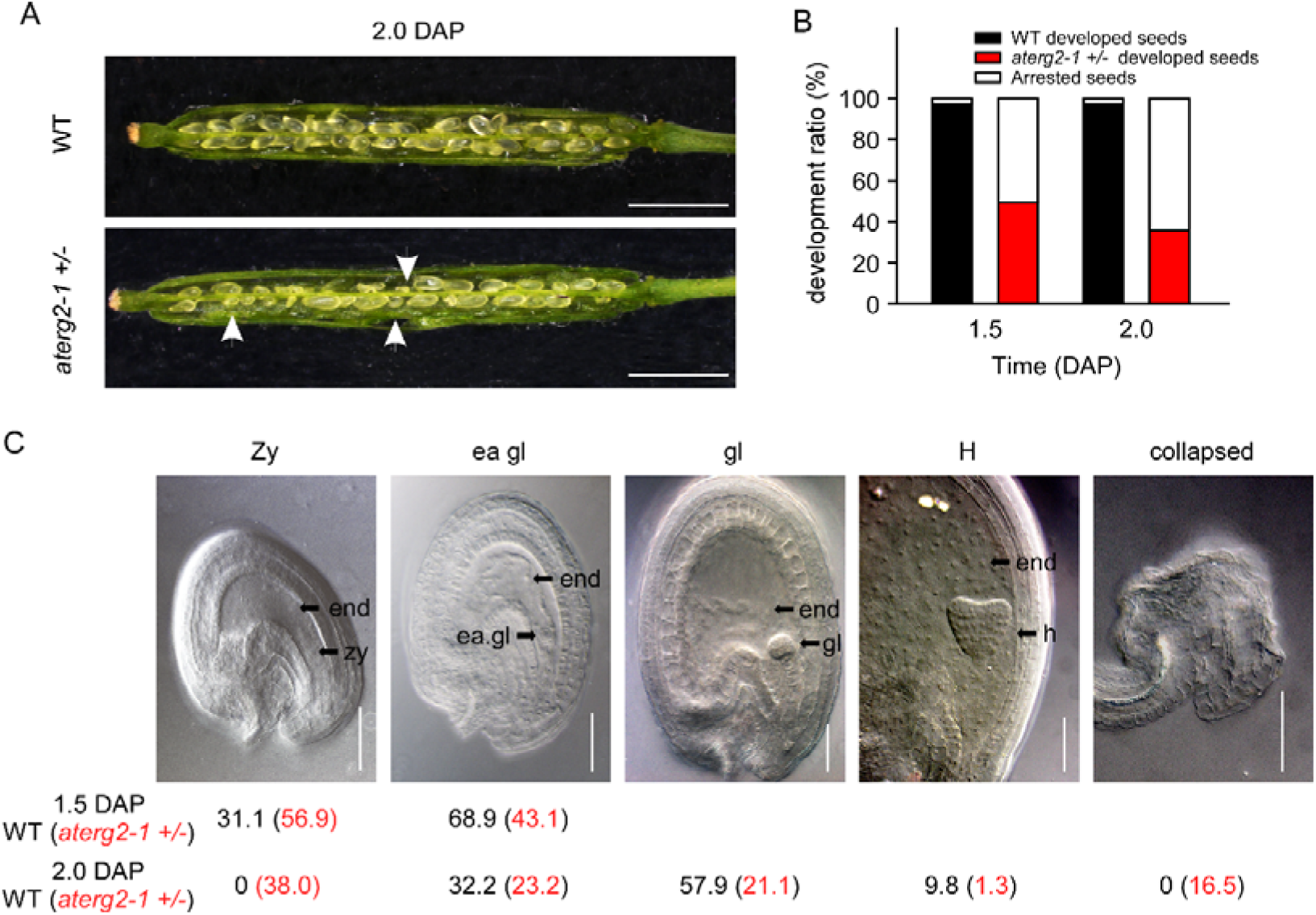
The development phenotypes of seeds and ovule in WT and *aterg2-1* +/- at 1.5 and 2.0 DAP. (A) The opened 2.0-DAP siliques from WT (top) and *aterg2-1* +/- (bottom). The arrows point to the aborted seeds. Bar = 1 mm. (B) The seed abortion ratio of WT and *aterg2-1* +/- at 1.5 and 2 DAP. Silique numbers = 8. (C) The ratio of developed ovules for WT and *aterg2-1* +/- at 1.5 and 2.0 DAP, respectively (n > 200). The embryos at the zygote stage are indicated by zy; the early globe-stage embryos (with fewer than four cells in a single embryo) are indicated by ea. gl; globe-stage embryos are indicated by gl; heart-stage embryos are indicated by H; the collapsed stage is indicated by the arrested seeds without the embryo. Bar = 0.1 mm

### ROS accumulation and activated transcription of ROS-responsible and cell death-related genes in *aterg2-1* +/- arrested seeds

Previous studies showed that the knockdown of ERAL1 in human cell lines by siRNA inhibits mitochondrial protein translation and elevates mitochondrial reactive oxygen species (ROS) production, leading to cell death (Uchiumi et al., 2010; Xie et al., 2012). Thus, ROS production and the expression of the ROS-related genes may cause the developmental stagnation of embryos and arrested seeds in *aterg2-1* +/-. To test this hypothesis, we performed DAB staining to detect ROS content in *Arabidopsis*. We found that the arrested seeds in *aterg2-1* +/- exhibited more pronounced increases in DAB staining than did the developed seeds in WT and *aterg2-1* +/- (Figure 5A), suggesting higher ROS levels in the arrested seeds of *aterg2-1* +/- than in the developed seeds of WT and *aterg2-1* +/-. Furthermore, *Alternative oxidase1a* (*AOX1a*), a nuclear gene encoding mitochondrial protein, is activated by ROS through ROS-responsible transcription factors WRKY40 and ANAC017 (Ng et al., 2013; Van Aken et al., 2013; Liu et al., 2014). Thus, these three genes can act as the marker genes for monitoring ROS levels. qRT-PCR showed that the transcription levels of these three genes was specifically higher at 1.5 and 2.0 DAP in the arrested seeds of *aterg2-1* +/- than in the developed seeds of WT and *aterg2-1* +/- (Figure 5B).

**Figure 5.**
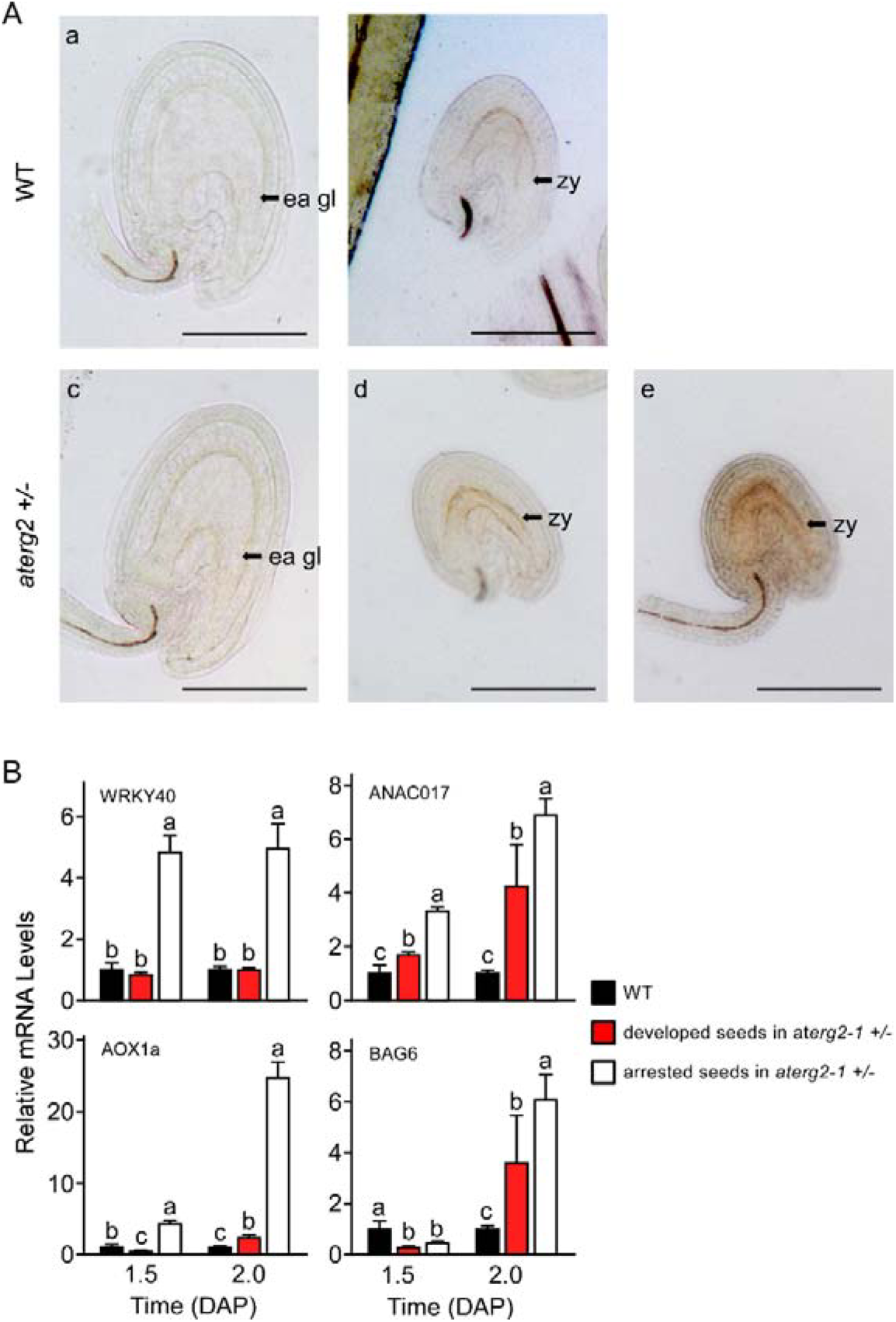
ROS accumulation and expression of ROS-responsible and cell death-related genes in the arrested seeds of *aterg2-1*. (A) DAB staining of 1.5-DAP seeds of WT and aterg2-1. The arrow points to the early globe-stage embryos (a and c) and the zygote embryos (b, d, and e). Bar=0.05 mm. (B) The relative expression levels of ROS-responsible genes *WRKY40*, *ANAC017*, and *AOX1a*, and cell death-related genes *AtBAG6* were determined using qRT-PCR in 1.5- and 2.0-DAP seeds of WT and *aterg2-1*. The mRNA levels were normalised to that of actin. The dark bars represent the WT seeds, and the red and white bars represent the developed and arrested seeds of *aterg2-1*, respectively. Error bars indicate the SD of three biological replicates. Different letters (a–c) indicate statistically significant differences (p < 0.05) according to Tukey’s multiple range test.

The <u>B</u>CL2-<u>a</u>ssociated athanogene (BAG) family, as the calmodulin-binding proteins, serve as co-chaperones to recruit HSP70/HSC70 to a specific target protein regulating diverse cellular pathways, such as programmed cell death (PCD) and stress responses in eukaryotes (Hung et al., 2003; Yan et al., 2003). Kang *et al* (Kang et al., 2006) showed that overexpression of *AtBAG6* induced cell death phenotypes consistent with programmed cell death (PCD) in yeast and plants. Here, we found that *AtBAG6* is extremely up-regulated in 2.0-DAP seeds of *aterg2-1* +/- but not in WT plants (Figure 5B), which suggested that inactivation of *AtERG2* induces the higher ROS levels and the cell death of embryos after pollination to produce the arrested seeds in *aterg2-1* +/- plants.

### Chloramphenicol treatment causes arrested development of the early embryo in *Arabidopsis*

To investigate whether the abortion of seeds in *aterg2-1* +/- is associated with mitochondria protein synthesis, we chose chloramphenicol (CAP), which prevents protein chain elongation by inhibiting the peptidyl transferase activity of the bacterial ribosome, to inhibit the translation of mitochondria-encoded genes and observe the phenotype of seed development in *Arabidopsis* WT plants. *Arabidopsis* flowers were treated with CAP solution before pollination until 2 DAPs using the floral dip method. We found that the silique length was shorter at 1.5 and 2.0 DAP with CAP treatment than in the control (Figure 6A), and the development rate of the embryo was 98.4% for the zygote period at 1.5 DAP and 62.5% for the zygote at 2.0 DAP with CAP treatment. However, the ratio of the zygote was 45.0% for 1.5 DAP and 8.2% for 2.0 DAP in the control (Figure 6B). Additionally, the transcription of *WRKY40*, *ANAC017*, *AOX1a*, and *AtBAG6* was induced with CAP treatment, and the transcription level of AtERG2 did not change (Figure 6C). These results indicated that CAP treatment plays a role similar to that of insertional inactivation of AtERG2 to trigger the ROS-related cell death through dysfunction of mitochondrial proteins.

**Figure 6.**
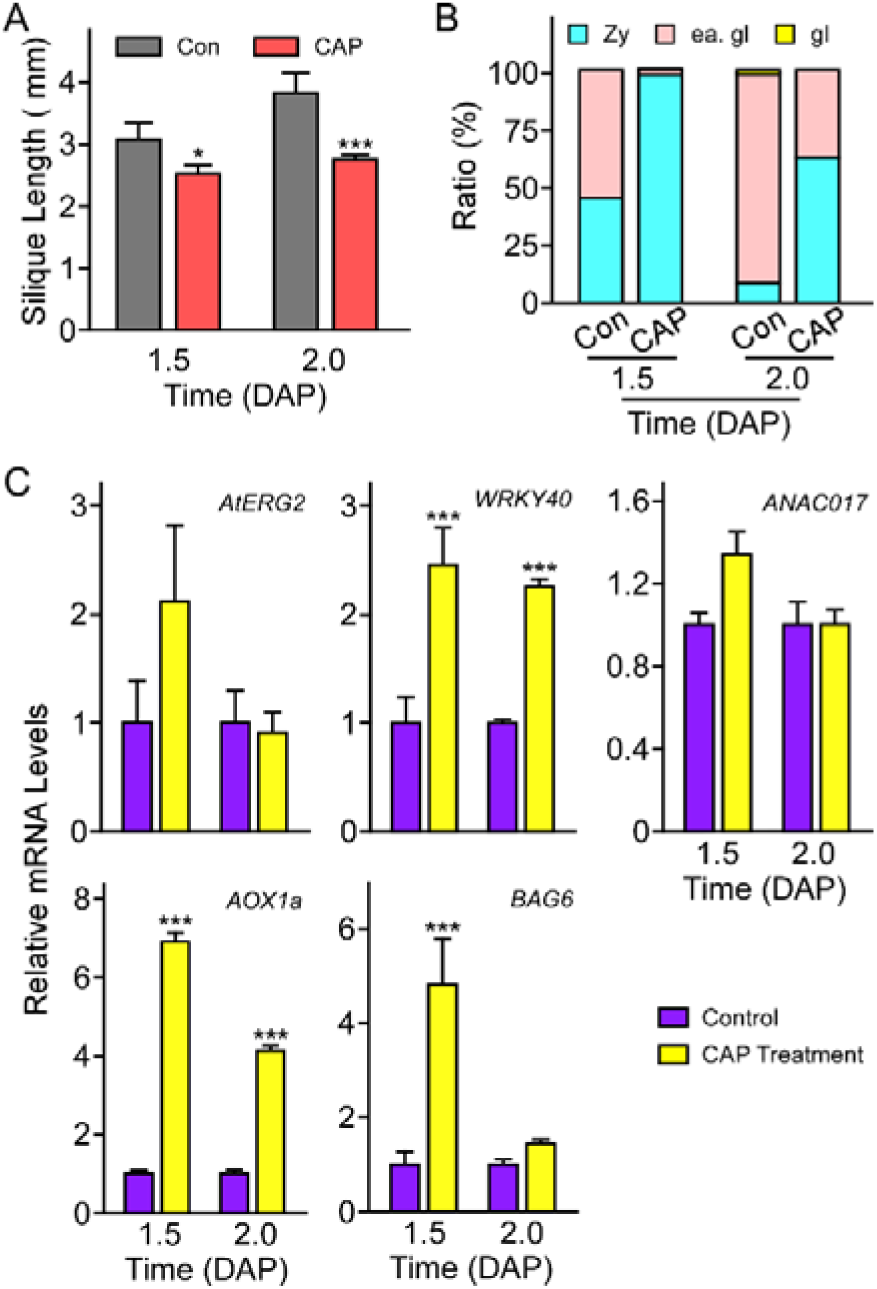
Chloramphenicol treatment causes seed abortion and cell death in *Arabidopsis*. (A) The length of 1.5- and 2.0-DAP siliques of wild-type (WT) plants after treatment with Chloramphenicol (CAP). The orange bars represent treatment with CAP solution (500 μg /mL CAP in 0.02% Silwet-77), the gray bars represent treatment with 0.02% Silwet-77 (Control). The values are means ± SD (n = 8). * indicates statistically significant values (P < 0.05) *** indicates extremely statistically significant values (P < 0.01). (B) The abortion ratio of 1.5- and 2.0-DAP seeds (n > 150) treated with CAP solution. The zygote embryo is indicated by zy; the early globular-stage embryo is indicated by ea gl; and the globular-stage embryo is indicated by gl. *** indicates extremely statistically significant values (P < 0.01). (C) The relative expression levels of AtERG2; ROS-responsible genes *WRKY40*, *ANAC017*, and *AOX1a*, and cell death-related genes *AtBAG6* were determined using qRT-PCR in 1.5- and 2.0-DAP seeds treated with CAP. The mRNA levels were normalised to that of *actin*. The yellow bars represent the seeds treated with CAP solution, and the purple bars represent the control. Error bars indicate the SD of three biological replicates. *** indicates extremely statistically significant differences (p < 0.01) according to Tukey’s multiple range test.

## Discussion

After double fertilisation in flowering plants, seed development starts with the coordinated development of embryo and endosperm, as well as interactions between endosperm and seed coat, which require both parental genomes. However, prior studies showed that embryo development is dominantly affected by both the female gametophyte and the sporophytic tissue of the parent plant, which is classified as the gametophytic maternal effect (GME) (Grossniklaus et al., 1998; Pien and Grossniklaus, 2007). Based on high-throughput sequencing, Autran *et al*. (Autran et al., 2011) showed that maternal epigenetic regulation controls the parental contributions in early embryogenesis in *Arabidopsis*. Additionally, mitochondria act as another maternal inheritance factor that affects the early seed development, although the ultrastructure observations implied that mitochondria could also be transmitted from sperm to the egg cell in tobacco (Yu and Russell, 1994). Here, we found that AtERG2 is a nucleus-encoded and mitochondria-localised ERA-like GTPase in *Arabidopsis*, and the T-DNA insertion mutant lines of *AtERG2* show a recessive lethal and GME phenotype after self-crossing or back-crossing with WT, which suggested that *aterg2* +/- represents a distinct class of GME mutants affecting early seed development.

The *Arabidopsis* genome contains three homologues of ERG: AtERG1 is localised in the chloroplast, AtERG2 is localised in mitochondria (Suwastika et al., 2014), and AtDER, similar to chloroplast-localised double Era-like GTPase in *Nicotiana benthamiana* (NbDER) (Jeon et al., 2014), is predicted to be localised in the chloroplast. In this study, we further showed that AtERG2 is localised to the mitochondria, dependent on its N-terminal sequence and is mainly expressed in the leaf vein, trichome, mature pollen, and ovule. The heterozygous *aterg2* +/- plant showed a severe abortion of seeds in siliques from 1.5 DAP, and more than 50% were arrested in an early stage of development, but only 2.4% were arrested in WT silique. After an additional 8 h of development, 16.5% of the seeds were severely crushed into a group of dead cell aggregates, and only 45.5% of embryos could successfully split out suspensors and develop into the globe stage. However, almost 100% of embryos developed into globe-stage embryos in WT silique. This phenomenon is similar to ERG function in *Antirrhinum majus* (Ingram et al., 1998), suggesting that mitochondria-localised *ERG* are essential for early seed development in plants.

The mitochondria genome size ranges from <6 kilobase pairs (kbp) in *Plasmodium falciparum* (the human malaria parasite) to >200 kbp in land plants (Gray et al., 1999), which encode a limited number of mitochondria tRNAs, ribosomal proteins, essential subunits of the different respiratory chain complexes, and some components involved in cytochrome-c biogenesis (Marienfeld et al., 1999). Therefore, mitochondria play a pivotal role in regulating cellular energy homeostasis and redox balance, and the disabled respiratory chain may cause broken electron transport chain balance and result in ROS accumulation in the matrix, possibly leading to cell death (Blackstone and Green, 1999). Here, we showed that AtERG2 binds mitochondrial ribosome small subunit component 18S rRNA, which benefits the maturation of 18S rRNA and the assembly of the mitochondrial ribosome. Therefore, disruption of AtERG2 in the *aterg2-1* +/- mutant affected the translation of the respiratory chain-related proteins to trigger ROS accumulation; it followed the expression of the ROS-related genes *WRKY40*, *ANAC017*, and *AOX1a* and the cell death-related gene BAG6, finally resulting in arrest of embryos at the zygote stage or early-globe stage in the aborted seeds of a mature silique from an *aterg2-1* +/- mutant. These results were further confirmed by CAP treatment of WT seeds because CAP caused the translational dysfunction of mitochondrial-encoded protein (Xie et al., 2012). These results are in accordance with previous studies, which showed that nuclear- or mitochondria-encoded gene mutations affecting mitochondrial functions, such as transcription, protein synthesis, and oxidative phosphorylation, affected ROS levels in developing gametophytes, thereby adversely affecting gametophyte and polar nuclei fusion during seed development (Maruyama et al., 2016; Pratibha et al., 2017). Our results suggest that AtERG2 promotes early seed development by affecting the maturation of the mitochondria ribosome small subunit and the mitochondrial protein translation in *Arabidopsis*, which highlights the role of mitochondrial ROS homeostasis in earlier seed development.

## Accession numbers

Sequence data for the genes in this study can be found in the GenBank database under the following accession numbers: AtERG2 (At1g30970), ANAC017 (AT1G34190), AOX1a (AT3G22370), WRKY40 (AT1G80840), BAG6 (AT2G46240), mitochondrial ribosome large subunit component 26S rRNA (AtMG00020), and small subunit 18S rRNA (AtMG01390).

## Funding

The work was supported by the National Natural Science Foundation of China (Grant No. 31070220) and the Specialized Research Fund for the Doctoral Program of Higher Education (20090003110016)

## Conflict of interest

The authors declare no conflict of interest.

## Supplementary Material

Table S1. **Primer sequences used in this study**.

Figure S1. **Phylogenetic analysis of AtERGs and homologous proteins**.

Phylogenetic tree constructed using the predicted amino acid sequences of the 21 ERA homologous proteins in 14 species. ERA (NP_417061.1) and EcDER (double Era-like GTPase, AAC75564) were from *E.coli*; ERAL1 (NP_005693.1) from *Homo sapiens*; DrERA (NP_001122219.1) from *Danio rerio*; MmERA (NP_071708.2) from *Mus musculus*; ScERA (NP_013736.1) from *Saccharomyces cerevisiae*; ERG (O82626.1) from *Antirrhinum majus*; AtERG1 (At5g66470), AtERG2 (At1g30960), and AtDER (At3g12080) from *Arabidopsis* thaliana; OsERG1 (Os05g49220), OsERG2 (Os08g10649.1), and OsDER (XP_015628913.1) from *Oryza sativa*; ZmERG (NP_001167959.1) and ZmDER (ACL53683) from *Zea mays*; StERG (XP_006363190.1) from *Solanum tuberosum*; CsERG (XP_004134545.1) from Cucumis sativus; NtERG (XP_016481550.1) from *Nicotiana tabacum*; SlERG (XP_004232641.1) from *Solanum lycopersicum*; NbDER (KC846070) from *Nicotiana benthamiana*.

Figure S2. **Schematic of different AtERG2-eYFP constructs**. The full-length and different truncated AtERG2 constructs were constructed with eYFP (green rectangles) attached to the C-terminal region. AtERG2 was divided into three different regions: 1–150 aa as the N-terminal of AtERG2 (AtERG2_N); 151–437 aa as the C-part of AtERG2 (AtERG2_C), which contains 151–326 aa as GTPase domain (AtERG2_GD); and 338–437 aa as the KH-domain (AtERG2_KH).

Figure S3. **Characterisation of T-DNA insertion mutant of *AtERG2***. (A) Two T-DNA insertion lines, *aterg2-1* (SALK_032115) and *aterg2-2* (SALK_032124), were identified by PCR. Wild-type (WT) *Arabidopsis* was used as the control. LP, LB, and RP primers are shown in Figure 1A and Table S1. (B) The mature seed numbers per silique of WT and mutant lines (n=60). (C) The ratio of developed and arrested seeds in WT and mutant lines at 6 DAP (number of siliques = 8).

Figure S4. **The ovule development in WT and *aterg2-1* +/- at Before pollination (BP) and 0.5 DAP**.

The ratio of developed ovules for WT and *aterg2-1* +/- at BP and 0.5 DAP, respectively (n > 150). The egg cell is indicated by ec; the synergid cells are indicated by sy; The two polar nuclei are indicated as pc; the central cell after the polar cell fused is indicated by cc; the early globe-stage embryos (with fewer than four cells in a single embryo) are indicated by ea. gl; the endosperm cell is indicated as end. Bar = 0.1 mm.

